# Broad-scale alpha- and beta- diversity patterns are robust to use of exact sequence variants versus operational taxonomic units

**DOI:** 10.1101/283283

**Authors:** Sydney I. Glassman, Jennifer B.H. Martiny

## Abstract

Recent controversy focuses on the best method for delineating microbial taxa, based on either traditional operational taxonomic units (OTUs) or exact sequence variants (ESVs) of marker gene sequences. We sought to test if the binning approach (ESVs versus OTUs defined by ≥97% sequence similarity) affected the conclusions of a large field study. The dataset included sequences targeting all bacteria (16S) and fungi (ITS), across multiple environments diverging markedly in abiotic conditions, over three collection times. Despite quantitative differences in microbial richness, we found that all alpha- and beta-diversity metrics were highly positively correlated (r > 0.90) between samples analyzed with both approaches. Consequently, statistical inferences were nearly indistinguishable. Further, ESVs only moderately increased the genetic resolution of fungal and bacterial diversity (1.3 and 2.1 times OTU richness, respectively). We conclude that for broad-scale (e.g., all bacteria or all fungi) alpha- and beta-diversity analyses, OTU or ESV methods will often reveal similar ecological results. Thus, while there are good reasons to employ ESVs, we need not question the validity of results based on OTUs. There are advantages and disadvantages of both, and which binning method to employ will depend on question/data in hand.

## Importance

Microbial ecologists have made exceptional improvements in our understanding of microbiomes in the last decade due to breakthroughs in sequencing technologies. These advances have wide ranging implications for fields ranging from agriculture to human health. Due to limitations in databases, the majority of microbial ecology studies use a binning approach to approximate taxonomy based on DNA sequence similarity. There remains extensive debate on the best way to bin and approximate this taxonomy. Here we examine two popular approaches using a large field based data set examining both bacteria and fungi, and conclude that there are not major differences in the ecological conclusions. Thus, despite controversy, it appears that standard microbial community analyses may not be very sensitive to the particulars of binning approaches.

## Main

Characterization of microbial communities by amplicon sequencing introduces biases and errors at every step. Hence, choices concerning all aspects of molecular processing from DNA extraction method (1) to sequencing platform (2) are contested. Further downstream, the options for computational processing of amplicon sequences are similarly deliberated (e.g. (3-5)). Yet despite these ongoing debates, microbial ecology has made great strides towards characterizing and testing hypotheses in environmental and host-associated microbiomes (e.g. (6, 7)).

Within microbiome studies, operational taxonomic units (OTUs) have been used to delineate microbial taxa, as microbial diversity still vastly outstrips our global databases (8). While any degree of sequence similarity could be used to denote individual taxa, a 97% sequence similarity cutoff became standard within microbial community analyses. This cutoff attempted to balance previous standards for defining microbial species (9) and a recognition of spurious diversity accumulated through PCR and sequencing errors (10, 11).

Recently, controversy over classifying microbial taxa has been renewed with the suggestion that taxa should be defined based on exact nucleotide sequences of marker genes.

Delineating taxa by exact sequence variants (ESVs), also termed amplicon sequence variants (ASVs)(12) or zOTUs (13), is not only expected to increase taxonomic resolution, but could also simplify comparisons across studies by eliminating the need for re-binning taxa when datasets are merged. Due to these advantages, there has been a surge in bioinformatic pipelines that seek to utilize ESVs and minimize specious sequence diversity (13-15). Moreover, proponents have stated that ESVs should replace OTUs altogether (12), and some journal reviewers are insisting that ESVs be used over OTUs as a condition for a study’s publication. However, as with the adoption of any new approach, there remains a need to quantify how the ESV method compares to the large body of previous research based on 97% OTUs. Further, OTU classifications remain biologically useful for comparing diversity across large datasets (7) or identifying clades that share traits (16).

Here, we tested if use of ESVs versus 97% OTUs affected the ecological conclusions, including treatment effects and alpha- and beta-diversity patterns, and if these conclusions varied based on the amplicon targeted. To do so, we used a dataset from a large leaf litter decomposition field study. This study included a “site” and “inoculum” treatment, in which all microbial communities were reciprocally transplanted into all five sites (see SI Methods) along an elevation gradient (17). We sequenced both bacteria (16S) and fungi (ITS2) from litterbags collected at three time points (6, 12, and 18 months after deployment) in separate Illumina MiSeq sequencing runs. While we expected that the binning approach would alter observed richness, we hypothesized that it might not alter trends in alpha- and beta-diversity, but that results might differ based on amplicon sequenced.

In total, we analyzed >15 million bacterial and >20 million fungal sequences using UPARSE v10 (**Table S1**), which allowed for a direct comparison of OTU versus ESV approaches by keeping all other aspects of quality filtering and merging consistent (4). ESV and OTU alpha-diversity was strongly correlated across samples using four metrics for both bacteria and fungi (mean Pearson r = 0.95 ± 0.02; all P values < 0.001). For the three metrics (Berger-Parker, Shannon, Simpson), the ESV and OTU approaches were not only highly correlated (mean Pearson r = 0.95 ± 0.02), but nearly equivalent in their values (mean slope = 0.97; **Table S2**). For observed richness, ESV versus OTU was also highly correlated across all time points/sequencing runs (Pearson r > 0.92; **Figure 1A,B**). However, bacterial OTU richness was approximately half that of ESV richness for the same sample (mean slope = 0.46), and fungal OTU richness was approximately three quarters that of ESV richness (mean slope = 0.79). We speculate that this difference between bacteria and fungi is due to the coarser phylogenetic breadth of the 16S versus ITS genetic regions.

**Figure 1.**
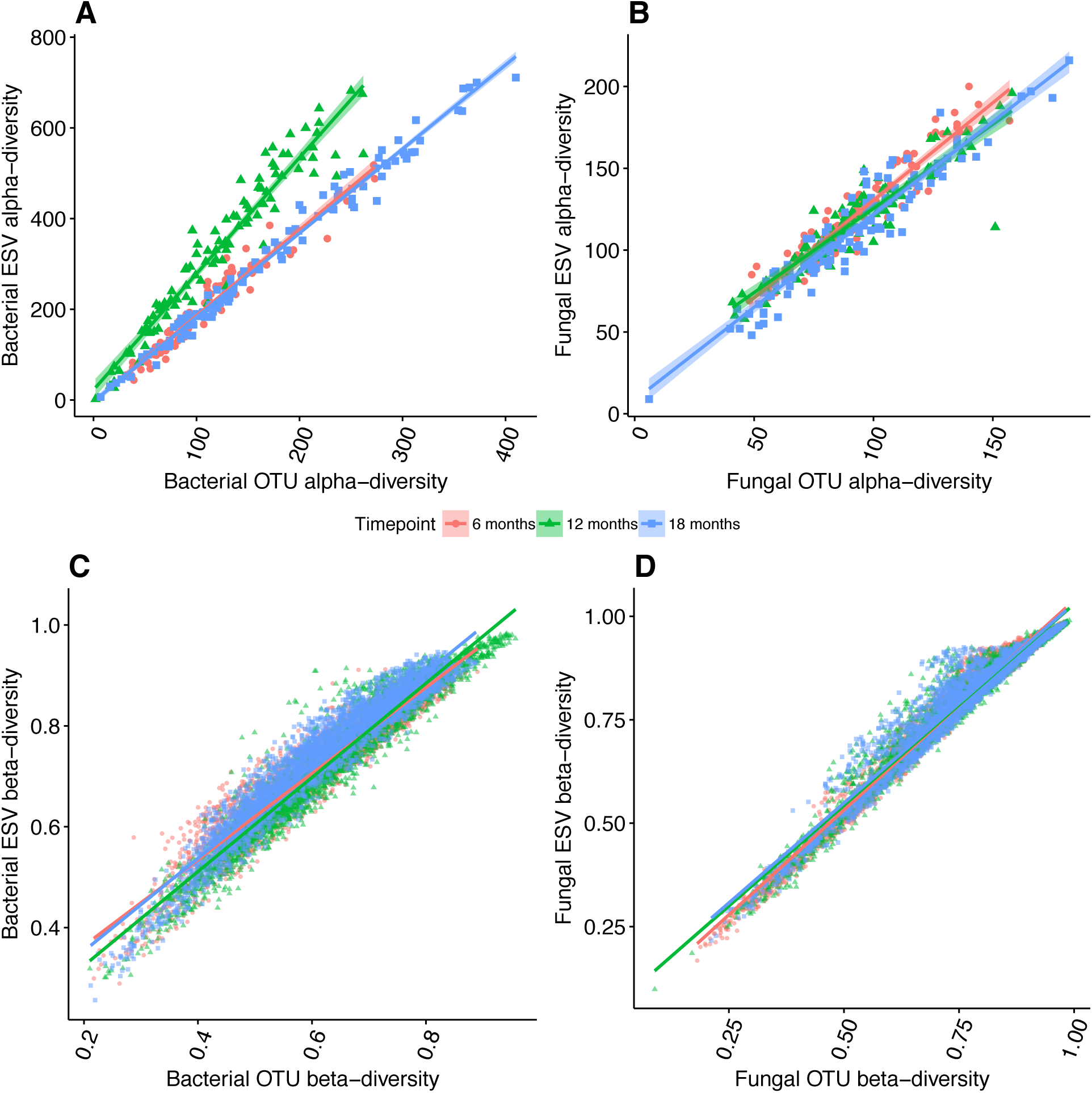
Comparison of observed alpha diversity for A) bacteria and B) fungi as assayed by the richness of 97% similar operational taxonomic units (OTU) versus exact sequence variants (ESV). Numbers are total observed richness after normalizing to 10,000 sequences per sample from three time points (16, 12, 18 months). Comparison of observed beta diversity for C) bacteria and D) fungi as assayed by the Bray-Curtis dissimilarity for OTUs versus ESVs from three time points (16, 12, 18 months).

Beta-diversity metrics were also strongly correlated across samples for ESVs and OTUs (Bray-Curtis average Mantel r = 0.96 for bacteria and 0.98 for fungi; all P < 0.01; **Figure 1C,D**), whether assessed by abundance-based (Bray-Curtis) or presence-absence (Jaccard) metrics (**Table S2)**. Moreover, the values of the beta-diversity metrics were nearly identical regardless of binning approach (slopes ~1).

The highly correlated alpha and beta-diversity metrics indicated that results based on these metrics should yield similar ecological conclusions. Indeed, the patterns of bacterial and fungal richness and community composition across the elevation gradient were nearly indistinguishable (**Figures 2 & S1**) as were the statistical tests for both richness (**Tables S3**) and community composition (**Tables S4, S5**). Moreover, family- and genus-level composition at each site along the gradient was virtually identical for bacteria **(Figure S2)** and highly similar for fungi (**Figure S3**). We also included a mock community of eight distinct bacterial species in our PCR and sequencing runs. Both approaches resulted in highly similar mock community composition (**Figure S4**). Thus, we found no evidence that ESVs yield better taxonomic resolution or are more sensitive to detecting treatment effects (12). If anything, the ESV method appeared to be slightly less sensitive to detecting treatment effects on richness than the OTU method, especially for fungi in which fewer significant treatment effects were detected using ESVs **(Table S3)**.

**Figure 2.**
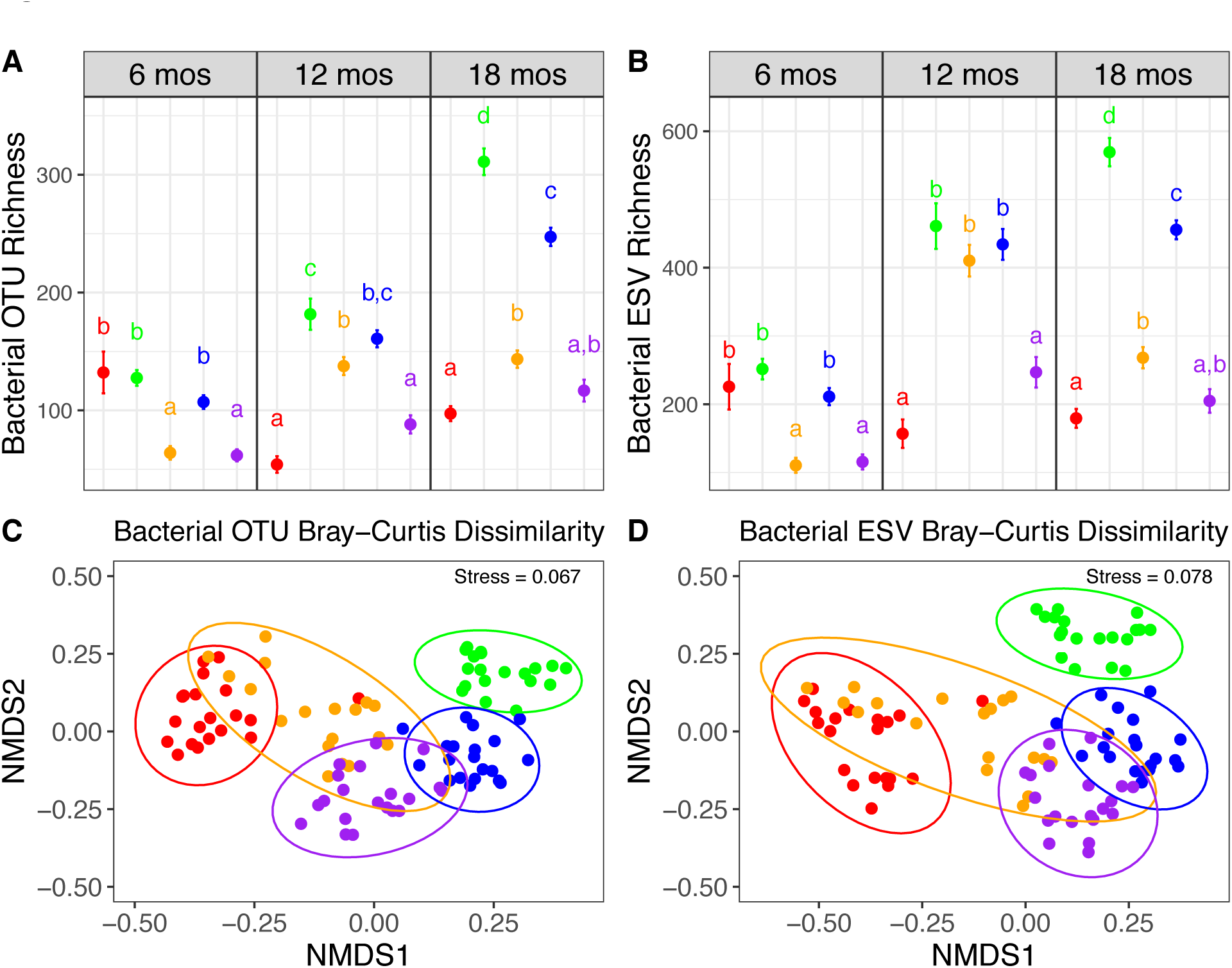
Comparison of alpha diversity results using A) operational taxonomic units (OTUs) versus B) exact sequencing variants (ESVs) for bacteria across elevation gradient at three time points (16, 12, 18 months). Each point represents mean observed richness per litterbag per site, and lines indicated standard error (averaged across five inoculum treatments and four replicates; n=20). Letters represent Tukey HSD significant differences across sites within a time point. Comparison of beta-diversity results using NMDS ordination of Bray-Curtis community dissimilarity of C) bacterial OTUs and D) bacterial ESVs colored by site at final time point (18 months). Ellipses represent 95% confidence intervals around the centroid. Colors represent sites along the elevation gradient ranging from lowest elevation (red; 275m) to highest elevation (purple 2240m) with middle elevation sites colored as follows: green = 470m; orange= 1280 m, blue = 1710 m).

Despite quantitative differences in microbial richness, ecological interpretation of our large bacterial and fungal community dataset was robust to the use of ESVs versus 97% OTUs. Thus, even though there are good reasons to take an ESV approach, we need not question the validity of studies based on 97% OTUs. Indeed, while previous studies have found that ESVs can help explain additional variation among samples (18, 19), the general alpha- and beta-diversity patterns of ESVs and OTUs in these studies were also quite similar. In general, we suspect that the robustness of such comparisons will vary depending on the breadth of the microbial community targeted. For instance, here we characterized all bacteria and fungi in a diverse environmental community, as opposed to a narrower subset of taxa or a less diverse host-associated community.

It is clear that both 97% OTUs and ESVs mask ecologically important trait variation of individual taxa (18, 20). In our study, ESVs only slightly increased the detection of fungal and bacterial diversity (1.3 and 2.1 times OTU richness, respectively), highlighting that ribosomal marker genes at any resolution are generally poor targets for improving genetic resolution within a microbial community. For example, it is widely known that many taxa can share the same 16S (20) or ITS (21). Thus, if strain identification is critical, then a full genome (22) or amplicon of a less conservative marker gene (23) is required. But for broad-scale community alpha and beta-diversity patterns, although the vagaries of molecular and bioinformatics processing inevitably add noise to microbial sequencing data, strong community-level signals will likely emerge with suitable study designs and statistics regardless of binning approach.

## Acknowledgments

We thank C.Weihe, J. Li, M.B.N Albright, C. Looby, A.C. Martiny, K.K. Treseder, S.D. Allison, M. Goulden, A.B. Chase, K.E. Walters, K. Isobe for their assistance in setting up the reciprocal transplant experiment and data collection used for this analysis. We thank A.A. Larkin, A.B. Chase, K.E. Walters, and K. Isobe for helpful comments on the manuscript. This work was supported by the National Science Foundation (DEB-1457160) and the U.S. Department of Energy, Office of Science, Office of Biological and Environmental Research under award DE-PS02-09ER09-25.

## Methods

The dataset is derived from a reciprocal transplant experiment conducted with leaf litter across five sites ranging in elevation from 275 to 2240m in southern California, USA (Glassman et al in prep). Precipitation and temperature co-vary along this elevation gradient (275m, 470m, 1280m, 1710m, 2240m). Total precipitation throughout the duration of the experiment ranged from 213 to 1415mm and mean soil temperature ranged from 11 to 26 °C. As part of the transplant, we constructed microbial litterbags with 0.22 µm nylon mesh that allow for movement of water and nutrients but prevent dispersal of microbes (1). We filled each bag with a common substrate of 5g of homogenized, gamma-irradiated, ground-up litter from a middle elevation site. We then inoculated each bag with 50mg of homogenized, ground-up litter containing the natural microbial community from each site for the initial “inoculum”. At each site along the gradient, we deployed the bags into 4 replicate plots in October 2015, and collected samples at 6, 12, and 18 months until April 2017. DNA was extracted from 100 litterbags (5 sites x 5 inoculum treatments x 4 plots) at each of the three time points (n=300) using the FastDNA Spin Kit for Soil (MP Biomedicals, Santa Ana, CA, USA) following the manual with the modification of adding three freeze/thaw cycles (30s in liquid Nitrogen followed by 3-5 min in 60°C water bath) prior to bead beating step to improve cell lysis.

To characterize bacterial composition, we amplified the V4 region of the 16S ribosomal RNA (rRNA) gene using the 515F-926R primers (2) with modifications to improve diversity (3) with the forward primer as the bar-coded primer. PCR mixtures for amplification contained 0.2 µL of NEB Hot Start Taq (5 units/µL) DNA polymerase (New England Biolabs, Ipswich, MA, USA), 2.5 µL of 10× 5Prime Hotmaster buffer minus MgCl_2_ (Quantabio, Beverly, MA, USA), 0.6 µL MgCl_2_ (50 mM), 0.50 µL dNTPs (10mM), 0.50 µL of 10 µM non-barcoded primer, 5 µL of 1 µM barcoded primer, 0.25 µL of BSA (20mg/ml), 5 µL of DNA template (diluted to 1:10 or 1:50 to overcome inhibitors), and water up to 25 µL. PCR conditions were as follows: denaturation at 94°C for 3 min; 35 amplification cycles of 45 s at 94°C, 30 s at 55°C, 20 s at 68°C, followed by a 10-min final extension at 68°C. For the 16S libraries, we included a mock community of eight bacteria strains from Zymo Microbiomics (Zymo Research, Irvine, CA, USA) that we PCR amplified and included in each sequencing run.

To characterize fungal composition, the ITS2 region of the Internal Transcribed Spacer (ITS) was amplified using the ITS9f-ITS4 primer combination designed by (4) and modified for Illumina MiSeq (5) following a staggered design (6). PCR mixtures for amplification contained 21.5 µl Platinum PCR Supermix (1.1x, Thermo Scientific, Waltham, MA, USA), 1 µl BSA (10 mg/ml, NEB, Ipswich, MA), 0.75 µl of both primers (10 µM, ITS9f and barcoded ITS4) and 1 µl of DNA template (diluted to 1:10) for a 25 µL reaction. PCR conditions were as follows: denaturation at 94°C for 5 min; 35 cycles of 45 s at 95°C, 1min at 50°C, 90 s at 72°C, followed by a 10-min final extension at 72°C.

PCR products were pooled visually according to intensity of bands based on electrophoresis gel images (scaled as weak, medium, or strong bands). Samples were pooled into six separate libraries (3 time points for each amplicon with 100-150 samples each). The pooled libraries were then purified according to the AMPure XP magnetic Bead protocol (Beckman Coulter Inc, Brea, CA, USA). For 16S, AMPure magnetic beads were used. For ITS2, we followed the same protocol but instead used a homemade solution of Sera-mag SpeedBeads (Fisher Scientific). The purified libraries were quality checked with an Agilent BioAnalyzer 2100 at the UCI Genomics High-Throughput Facility (UC Irvine, CA, USA) for size and concentration. The libraries were then sequenced in six separate Illumina MiSeq PE runs (2 × 250 bp) at the DNA Technologies Core, UC Davis Genome Center, Davis, CA, USA. Sequences were submitted to the National Center for Biotechnology Information Sequence Read Archive under accession number SRPXXXXX.

All bioinformatics processing was conducted in UPARSE (7) version 10 (https://www.drive5.com/usearch/manual/uparse_pipeline.html). We processed each amplicon library by each timepoint in order to examine variation in patterns among sequencing runs. We chose the UPARSE pipeline for ease of comparison of OTU versus ESV methods while keeping all other aspects of quality filtering and merging the same (7). First, primers were stripped, then reads were truncated based on quality, forward and reverse reads were merged, and then merged pairs were quality filtered using the fastq_filter command with a fastq_maxee parameter of 1.0. Next, at the same point in the pipeline, UPARSE can process both 97% OTUs with the “-cluster_otus” command and ESVs with the “unoise3” command. We used the default settings for each function, which for OTUs removes singletons with the “minsize 2” parameter, and the default minimum of 8 sequences per cluster for “unoise3”. Both of these methods are open reference and thus capture novel diversity. OTU tables were made with the otutab command for both 97% OTUs and ESVs.

Taxonomy was assigned in QIIME 1.9 (8) using the assign_taxonomy.py command. For 16S, assignments were made with the Greengenes database (9) using rdp classifier and 0.80 similarity cutoff. For ITS2, we used the UNITE database (10), accessed on June 28, 2017, using blast and minimum E value of 0.001. For ITS2, only reads mapping to kingdom Fungi were retained, and for 16S, all reads mapping to chloroplasts, mitochondria, or unclassified were removed.

Preliminary alpha diversity analyses were conducted in UPARSE. Samples were normalized to 10,000 sequences per sample using the otutab_norm command and alpha diversity metrics were calculated on the subsampled OTU tables using the alpha_div command. Next, these richness metrics were imported to R (11), where we performed Pearson correlations and linear regressions to determine the correlation, intercept, and slope of the relationship between four separate alpha diversity metrics (Berger-Parker, Observed Richness, Shannon, Simpson) for both the OTU and ESV approaches for each amplicon and at each time point. We then performed ANOVA and post-hoc Tukey HSD tests to determine if the significance of our treatments on observed richness were different for OTU versus ESV for each time point for each amplicon. All figures were made using the ggplot2 package (12) in the R software environment.

For beta diversity analyses, we took the raw OTU and ESV tables from UPARSE and calculated Bray-Curtis and Jaccard dissimilarity matrices for bacteria and fungi at each time point in the R package vegan (13) using the avgdist function (https://github.com/vegandevs/vegan/blob/master/man/avgdist.Rd). Specifically, we used this function to calculate a median, square-root transformed, Bray-Curtis or Jaccard dissimilarity matrix based on 100 subsamples of either 7,000 seq/sample for bacteria or 17,000 seq/sample for fungi. We then ran Mantel correlations in vegan to test the correlation between beta-diversity metrics between the binning methods for bacteria in fungi. We visualized these correlation in ggplot. Next, PERMANOVA tests were conducted with the Adonis function in vegan to test the effects of our two treatments and their interactions on the Bray-Curtis community dissimilarity of both bacteria and fungi as assessed by either OTUs or ESVs. We then visualized these patterns with NMDS using the metaMDS function in vegan.

To determine if binning method (OTUs vs ESVs) affected distribution of taxonomic groups among each site, we summarized the top 12 most abundant families or genera at each site for the inoculum leaf litter using the dplyr package and visualized this information with barplots the ggplot2 package in R. All data and scripts to re-create all figures and statistics from paper can be found on github: https://github.com/sydneyg/OTUvESV.

## Supplemental Information Figures S1-S4

**Figure S1.**
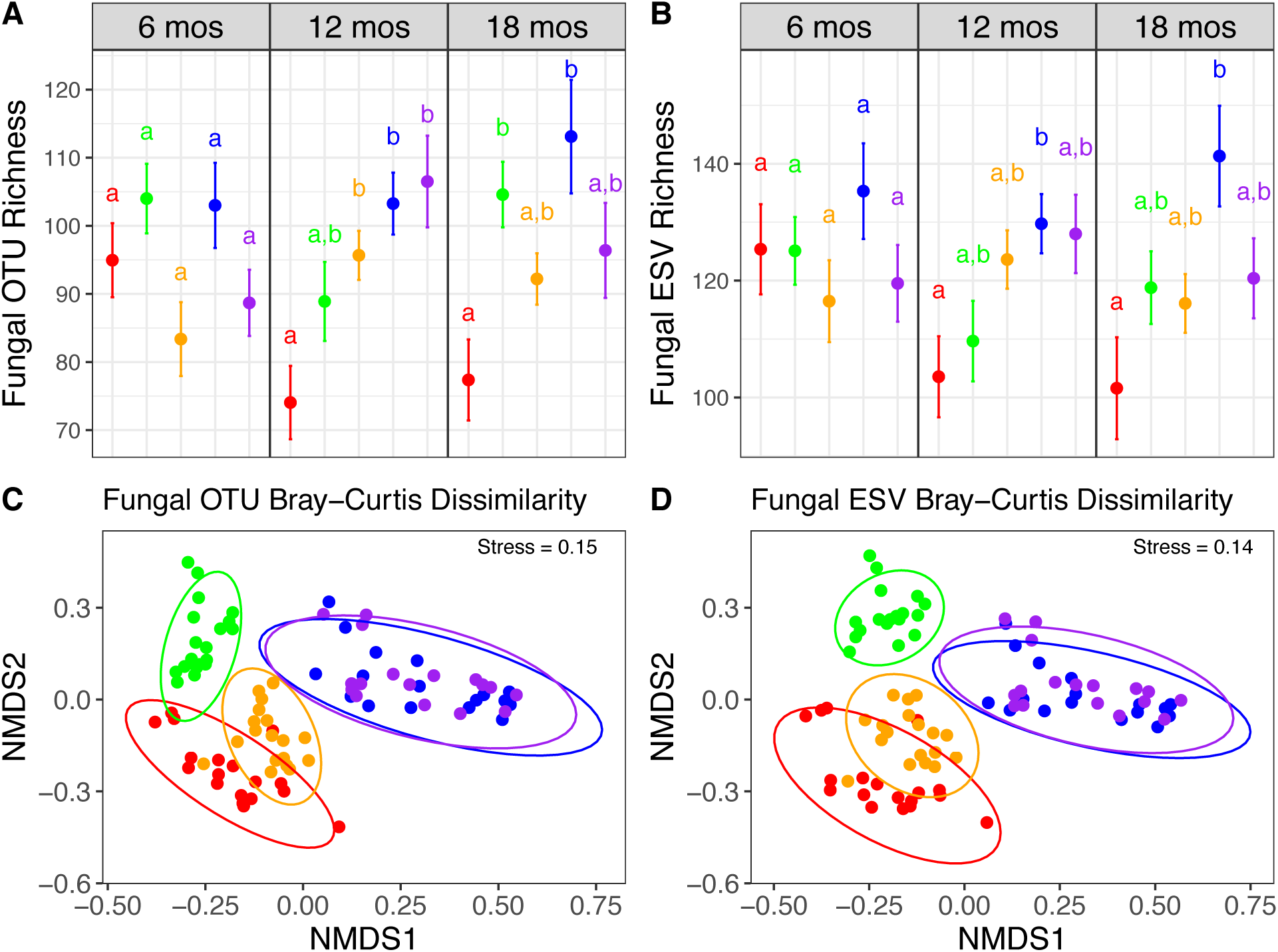
Comparison of alpha diversity results using A) operational taxonomic units (OTUs) versus B) exact sequencing variants (ESVs) for fungi across elevation gradient at three time points (16, 12, 18 months). Each point represents mean richness per litterbag per site (averaged across five inoculum treatments and four replicates; n=20). Letters represent Tukey HSD significant differences across sites within a time point. Comparison of beta-diversity results using NMDS ordination of Bray-Curtis community dissimilarity of A) fungal OTUs and b) fungal ESVs colored by inoculum at final time point (18 months). Ellipses represent 95% confidence intervals around the centroid. Colors represent inoculum from sites along the elevation gradient ranging from lowest elevation (red; 275m) to highest elevation (purple 2250m).

**Figure S2.**
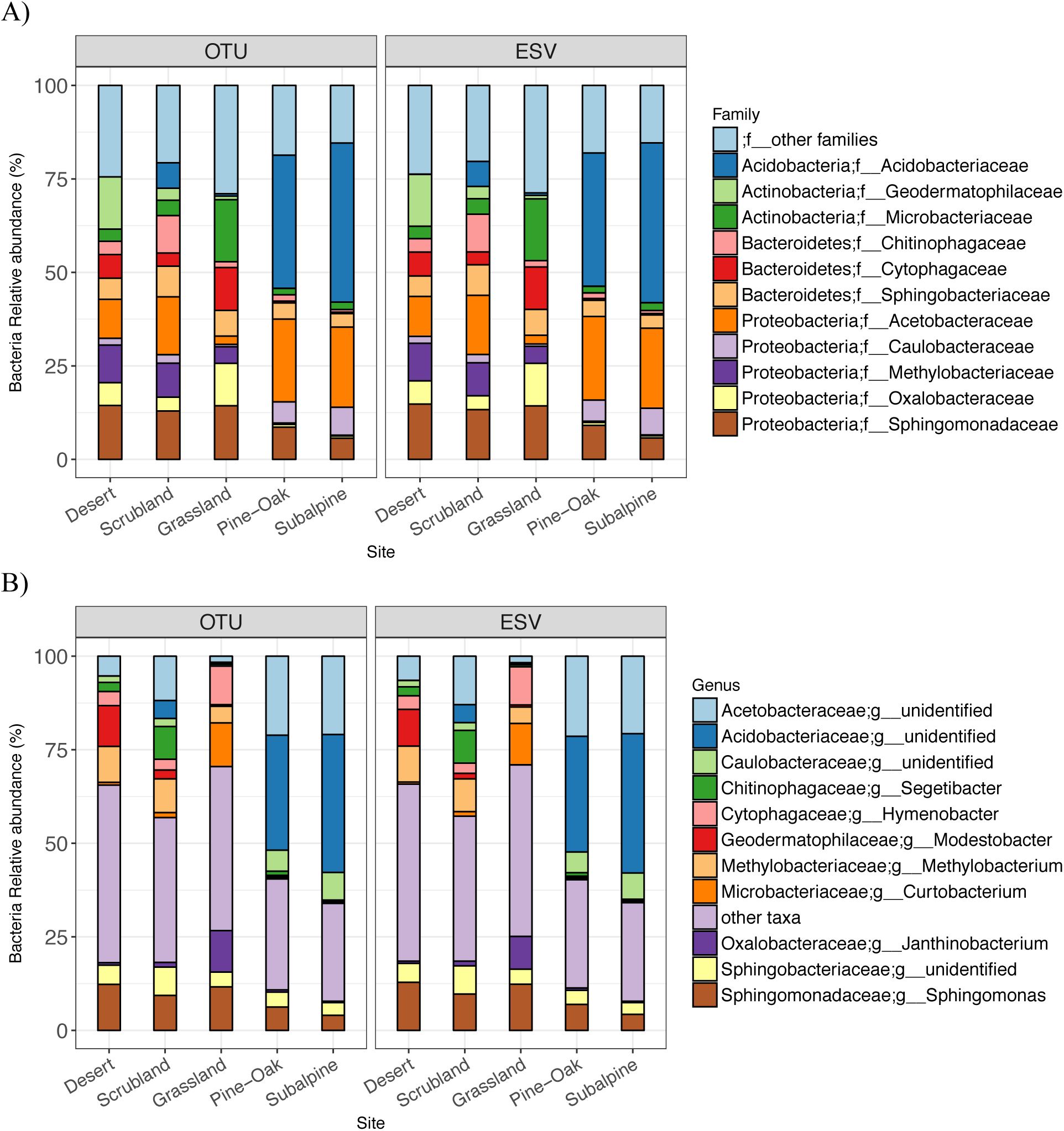
Relative abundance of bacterial sequences per sample of the inoculum leaf litter from each site (n=20) for OTUs versus ESVs summarized by A) the 12 most abundant families or B) the 12 most abundant genera.

**Figure S3.**
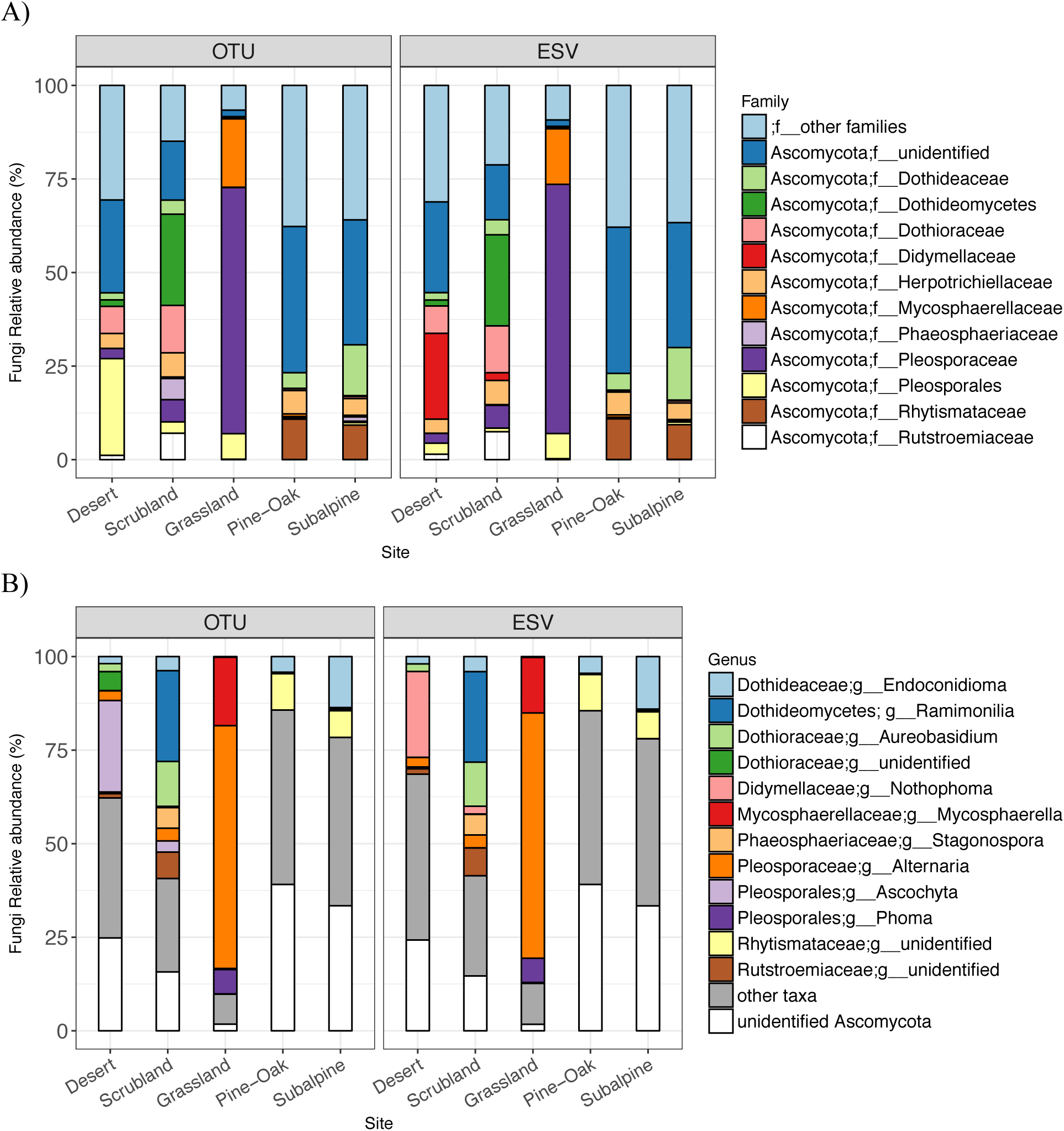
Relative abundance of fungal sequences per sample of the inoculum leaf litter from each site (n=20) for OTUs versus ESVs summarized by A) the 12 most abundant families (or next best taxonomic classification) or B) the 12 most abundant genera.

**Figure S4.**
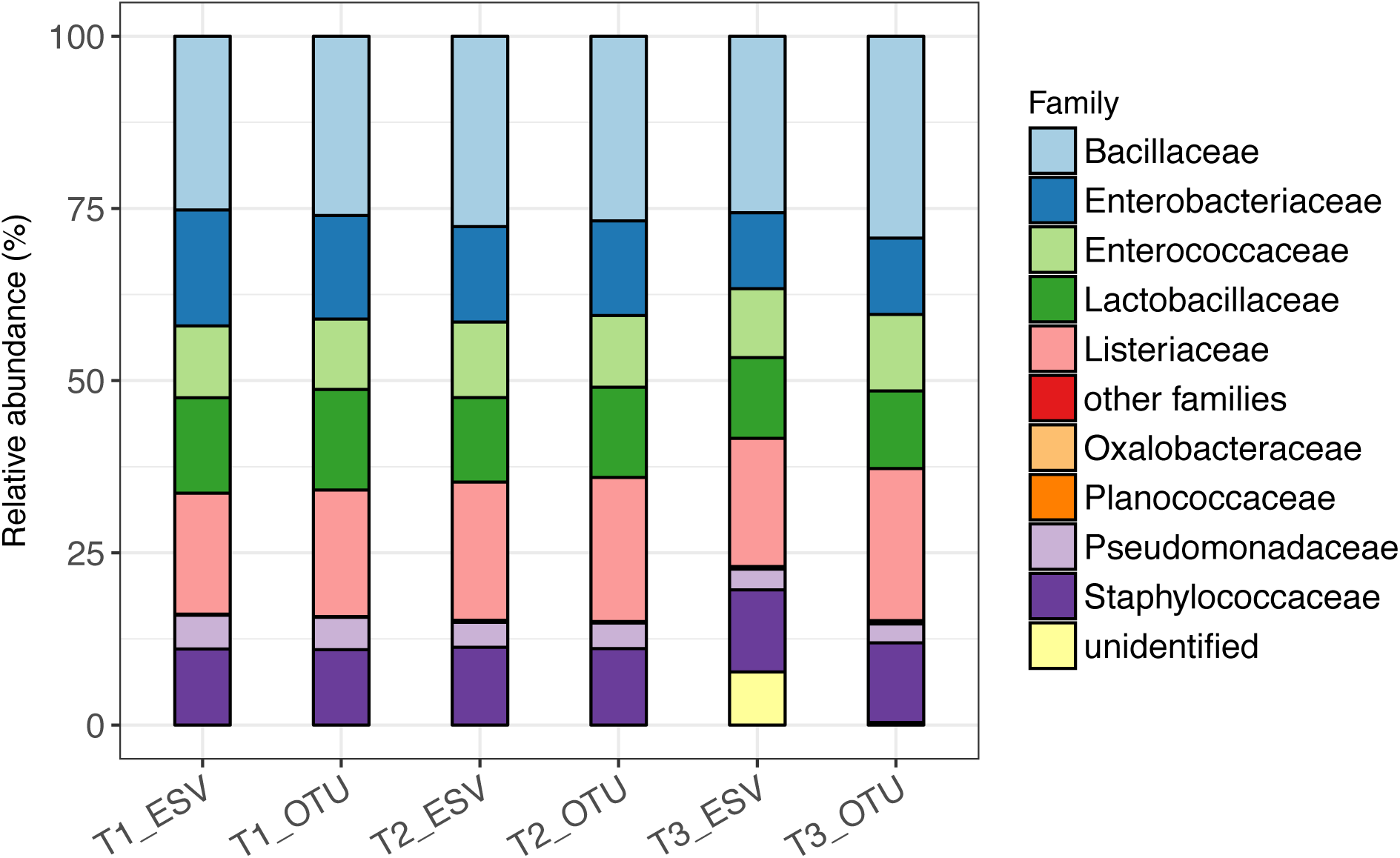
Relative abundance of bacterial sequences per mock community sample from each Illumina MiSeq run for each time point with either ESVs or 97% OTUs. The three timepoints represent 6 months (T1), 12 months (T2) and 18 months (T3) after transplantation. Both ESVs and OTUs largely recapitulated the eight mock community taxa at the family level, although in both cases more than 8 taxa were found. Bacterial taxa included in th mock community were: *Bacillus subtilis (F. Bacillaceae), Escherichia coli (F. Enterobacteriaceae), Salmonella enterica (F. Enterobacteriaceae), Enterococcus faecalis (F. Enterococcaceae), Lactobacillus fermentum (F. Lactobacillaceae), Pseudomonas aeruginosa (F. Pseudomonadaceae;), Staphylococcus aureus (F. Staphylococcaceae), Listeria monocytogenes (F. Listeriaceae).* For ESVs, there were more taxa within each of these dominant families than for OTUs, but since all the taxa included in the mock community have 16S multiple copies, it is unclear if these are truly spurious taxa or real genetic variation within the 16S gene (for information on taxa in mock community see: https://d2gsy6rsbfrvyb.cloudfront.net/media/amasty/amfile/attach/_D6300_ZymoBIOMICS_Microbial_Community_Standard_v1.1.3.pdf). For both OTUs and ESVs, there were low abundance sequences from families common in the experimental samples (i.e. Oxalobacteraceae), and these likely represent spillover between the barcoded samples.

## Supplemental Information Tables S1-S5

**Table S1.** Summary of data on Illumina MiSeq runs. Number of reads from each of the three different runs (corresponding to the three time points 6, 12, and 18 months), number of samples, and the number of OTUs versus ESVs for fungal and bacterial amplicons.

| Time Point | Amplicon | Kingdom | No. reads after quality filtering | No. Samples | No. OTUs | No. ESVs |
| --- | --- | --- | --- | --- | --- | --- |
| 6 months | ITS2 | Fungi | 5.9M | 95 | 1196 | 1240 |
| 12 months | ITS2 | Fungi | 7.3M | 95 | 1420 | 1594 |
| 18 months | ITS2 | Fungi | 7.3M | 114 | 1313 | 1243 |
| 6 months | 16S | Bacteria | 3.99M | 97 | 745 | 1306 |
| 12 months | 16S | Bacteria | 5.75M | 102 | 1163 | 1999 |
| 18 months | 16S | Bacteria | 5.4M | 119 | 1999 | 2513 |

**Table S2.** Pearson correlations and linear model relationships for 97% OTUs vs ESVs for alpha-diversity and beta-diversity metrics for bacteria (16S) and fungi (ITS2) by time point.

| Alpha-Diversity |  |  | Pearson correlation |  | Linear model |  |  |
| --- | --- | --- | --- | --- | --- | --- | --- |
| Microbe | Timepoint | Metric | r | P | Slope | R<sup>2</sup> | P |
| Bacteria | 6 mos | Berger-Parker | 0.95 | <0.001 | 0.97 | 0.9 | <0.001 |
| Bacteria | 12 mos | Berger-Parker | 0.92 | <0.001 | 0.97 | 0.84 | <0.001 |
| Bacteria | 18 mos | Berger-Parker | 0.95 | <0.001 | 1.06 | 0.91 | <0.001 |
| Fungi | 6 mos | Berger-Parker | 0.94 | <0.001 | 0.97 | 0.89 | <0.001 |
| Fungi | 12 mos | Berger-Parker | 0.92 | <0.001 | 0.94 | 0.84 | <0.001 |
| Fungi | 18 mos | Berger-Parker | 0.97 | <0.001 | 0.94 | 0.94 | <0.001 |
| Bacteria | 6 mos | Richness | 0.97 | <0.001 | 0.5 | 0.94 | <0.001 |
| Bacteria | 12 mos | Richness | 0.96 | <0.001 | 0.36 | 0.93 | <0.001 |
| Bacteria | 18 mos | Richness | 0.99 | <0.001 | 0.53 | 0.99 | <0.001 |
| Fungi | 6 mos | Richness | 0.95 | <0.001 | 0.75 | 0.9 | <0.001 |
| Fungi | 12 mos | Richness | 0.92 | <0.001 | 0.82 | 0.85 | <0.001 |
| Fungi | 18 mos | Richness | 0.95 | <0.001 | 0.8 | 0.91 | <0.001 |
| Bacteria | 6 mos | Shannon | 0.98 | <0.001 | 0.86 | 0.97 | <0.001 |
| Bacteria | 12 mos | Shannon | 0.95 | <0.001 | 0.77 | 0.9 | <0.001 |
| Bacteria | 18 mos | Shannon | 0.99 | <0.001 | 0.92 | 0.97 | <0.001 |
| Fungi | 6 mos | Shannon | 0.95 | <0.001 | 0.96 | 0.91 | <0.001 |
| Fungi | 12 mos | Shannon | 0.95 | <0.001 | 0.9 | 0.89 | <0.001 |
| Fungi | 18 mos | Shannon | 0.98 | <0.001 | 0.96 | 0.95 | <0.001 |
| Bacteria | 6 mos | Simpson | 0.96 | <0.001 | 1.01 | 0.92 | <0.001 |
| Bacteria | 12 mos | Simpson | 0.93 | <0.001 | 1.07 | 0.87 | <0.001 |
| Bacteria | 18 mos | Simpson | 0.95 | <0.001 | 1.11 | 0.91 | <0.001 |
| Fungi | 6 mos | Simpson | 0.95 | <0.001 | 1.07 | 0.9 | <0.001 |
| Fungi | 12 mos | Simpson | 0.93 | <0.001 | 0.98 | 0.86 | <0.001 |
| Fungi | 18 mos | Simpson | 0.98 | <0.001 | 1 | 0.96 | <0.001 |
| Beta-Diversity |  |  | Mantel correlation |  | Linear model |  |  |
| Microbe | Timepoint | Metric | Mantel r | P.value | Slope | R<sup>2</sup> | P |
| Bacteria | 6 mos | Bray-Curtis | 0.95 | 0.001 | 0.85 | 0.9 | <0.001 |
| Bacteria | 12 mos | Bray-Curtis | 0.96 | 0.001 | 0.93 | 0.93 | <0.001 |
| Bacteria | 18 mos | Bray-Curtis | 0.96 | 0.001 | 0.93 | 0.93 | <0.001 |
| Fungi | 6 mos | Bray-Curtis | 0.99 | 0.001 | 1.02 | 0.98 | <0.001 |
| Fungi | 12 mos | Bray-Curtis | 0.98 | 0.001 | 0.98 | 0.96 | <0.001 |
| Fungi | 18 mos | Bray-Curtis | 0.97 | 0.001 | 0.97 | 0.95 | <0.001 |
| Bacteria | 6 mos | Jaccard | 0.93 | 0.001 | 0.88 | 0.87 | <0.001 |
| Bacteria | 12 mos | Jaccard | 0.94 | 0.001 | 0.93 | 0.88 | <0.001 |
| Bacteria | 18 mos | Jaccard | 0.96 | 0.001 | 0.84 | 0.92 | <0.001 |
| Fungi | 6 mos | Jaccard | 0.99 | 0.001 | 1.1 | 0.97 | <0.001 |
| Fungi | 12 mos | Jaccard | 0.98 | 0.001 | 1.04 | 0.97 | <0.001 |
| Fungi | 18 mos | Jaccard | 0.98 | 0.001 | 1.06 | 0.97 | <0.001 |

**Table S3.** ANOVA results for OTUs and ESVs at each of the three time points, testing for effects of Site, Inoculum, and their interaction on bacterial or fungal richness.

| **Bacteria** | **6 months** | **OTU** |  |  |  |  |
| --- | --- | --- | --- | --- | --- | --- |
|  | Df | Sum Sq | Mean Sq | Fvalue | Pr(>F) |  |
| Site | 4 | 87281 | 21820 | 20.898 | 2.28E-11 | *** |
| Inoculum | 4 | 9952 | 2488 | 2.383 | 5.96E-02 | . |
| Site:Inoculum | 16 | 71114 | 4445 | 4.257 | 1.11E-05 | *** |
| Residuals | 70 | 73089 | 1044 |  |  |  |
| **Bacteria** | **6 months** | **ESV** |  |  |  |  |
|  | Df | Sum Sq | Mean Sq | F value | Pr(>F) |  |
| Site | 4 | 325282 | 81321 | 20.682 | 2.76E-11 | *** |
| Inoculum | 4 | 31158 | 7789 | 1.98E+00 | 1.07E-01 |  |
| Site:Inoculum | 16 | 286072 | 17880 | 4.547 | 4.33E-06 | *** |
| Residuals | 70 | 275243 | 3932 |  |  |  |
| **Bacteria** | **12 months** | **OTU** |  |  |  |  |
|  | Df | Sum Sq | Mean Sq | Fvalue | Pr(>F) |  |
| Site | 4 | 220757 | 55189 | 45.517 | < 2e-16 | *** |
| Inoculum | 4 | 25461 | 6365 | 5.25 | 0.000879 | *** |
| Site:Inoculum | 16 | 33138 | 2071 | 1.708 | 0.063351 | . |
| Residuals | 75 | 90937 | 1212 |  |  |  |
| **Bacteria** | **12 months** | **ESV** |  |  |  |  |
|  | Df | Sum Sq | Mean Sq | Fvalue | Pr(>F) |  |
| Site | 4 | 1414191 | 353548 | 37.559 | < 2e-16 | *** |
| Inoculum | 4 | 202288 | 50572 | 5.372 | 0.000737 | *** |
| Site:Inoculum | 16 | 268183 | 16761 | 1.781 | 0.049934 | * |
| Residuals | 75 | 705990 | 9413 |  |  |  |
| **Bacteria** | **18 months** | **OTU** |  |  |  |  |
|  | Df | Sum Sq | Mean Sq |  | Pr(>F) |  |
| Site | 4 | 660280 | 165070 | 134.294 | < 2e-16 | *** |
| Inoculum | 4 | 20634 | 5159 | 4.197 | 0.00407 | ** |
| Site:Inoculum | 16 | 24418 | 1526 | 1.242 | 2.59E-01 |  |
| Residuals | 74 | 90958 | 1229 |  |  |  |
| **Bacteria** | **18 months** | **ESV** |  |  |  |  |
|  | Df | Sum Sq | Mean Sq | Fvalue | Pr(>F) |  |
| Site | 4 | 2246528 | 561632 | 118.012 | <2e-16 | *** |
| Inoculum | 4 | 53361 | 13340 | 2.803 | 0.0317 | * |
| Site:Inoculum | 16 | 93870 | 5867 | 1.233 | 2.65E-01 |  |
| Residuals | 74 | 352173 | 4759 |  |  |  |
| **Fungi** | **6 months** | **OTU** |  |  |  |  |
|  | Df | Sum Sq | Mean Sq | Fvalue | Pr(>F) |  |
| Site | 4 | 6163 | 1540.9 | 4.265 | 0.00377 | ** |
| Inoculum | 4 | 12633 | 3158.3 | 8.741 | 8.53E-06 | *** |
| Site:Inoculum | 16 | 13216 | 826 | 2.286 | 0.00926 | ** |
| Residuals | 71 | 25654 | 361.3 |  |  |  |
| **Fungi** | **6 months** | **ESV** |  |  |  |  |
|  | Df | Sum Sq | Mean Sq | F value | Pr(>F) |  |
| Site | 4 | 3936 | 984 | 1.763 | 0.14594 |  |
| Inoculum | 4 | 24459 | 6115 | 1.10E+01 | 5.66E-07 | *** |
| Site:Inoculum | 16 | 23398 | 1462 | 2.62 | 0.00289 | ** |
| Residuals | 71 | 39636 | 558 |  |  |  |
| **Fungi** | **12 months** | **OTU** |  |  |  |  |

|  | Df | Sum Sq | Mean Sq | Fvalue | Pr(>F) |  |
| --- | --- | --- | --- | --- | --- | --- |
| Site | 4 | 12921 | 3230 | 7.075 | 7.64E-05 | *** |
| Inoculum | 4 | 7172 | 1793 | 3.927 | 0.00621 | ** |
| Site:Inoculum | 16 | 9612 | 601 | 1.316 | 0.21263 |  |
| Residuals | 70 | 31959 | 457 |  |  |  |

| **Fungi** | **12 months** | **ESV** |  |  |  |  |
| --- | --- | --- | --- | --- | --- | --- |
|  | Df | Sum Sq | Mean Sq | Fvalue | Pr(>F) |  |
| Site | 4 | 10530 | 2632.6 | 3.959 | 0.00593 | ** |
| Inoculum | 4 | 6069 | 1517.3 | 2.282 | 0.06907 | . |
| Site:Inoculum | 16 | 13794 | 862.1 | 1.296 | 0.22435 |  |
| Residuals | 70 | 46550 | 665 |  |  |  |

| **Fungi** | **18 months** | **OTU** |  |  |  |  |
| --- | --- | --- | --- | --- | --- | --- |
|  | Df | Sum Sq | Mean Sq |  | Pr(>F) |  |
| Site | 4 | 13800 | 3450 | 8.898 | 7.22E-06 | *** |
| Inoculum | 4 | 9753 | 2438 | 6.289 | 0.000221 | *** |
| Site:Inoculum | 16 | 27154 | 1697 | 4.377 | 7.50E-06 | *** |
| Residuals | 70 | 27141 | 388 |  |  |  |

| **Fungi** | **18 months** | **ESV** |  |  |  |  |
| --- | --- | --- | --- | --- | --- | --- |
|  | Df | Sum Sq | Mean Sq | Fvalue | Pr(>F) |  |
| Site | 4 | 15399 | 3850 | 6.472 | 0.000172 | *** |
| Inoculum | 4 | 10599 | 2650 | 4.455 | 0.002894 | ** |
| Site:Inoculum | 16 | 36416 | 2276 | 3.826 | 4.63E-05 | *** |
| Residuals | 70 | 41636 | 595 |  |  |  |

**Table S4.** PERMANOVA results for Bacterial OTUs vs ESVs, testing for effects of Site, Inoculum, and their interaction on Bray-Curtis community dissimilarity (data plotted in Figure 2C,D).

| **6 months** |  | OTU |  |  |  |  |  |
| --- | --- | --- | --- | --- | --- | --- | --- |
|  | Df | SumsOfSqs | MeanSqs | F.Model | R2 | Pr(>F) |  |
| Site | 4 | 5.3289 | 1.33222 | 17.4236 | 0.34467 | 1.00E-04 | *** |
| Inoculum | 4 | 2.5453 | 0.63633 | 8.3224 | 0.16463 | 1.00E-04 | *** |
| Site:Inoculum | 16 | 2.9225 | 0.18266 | 2.3889 | 0.18903 | 1.00E-04 | *** |
| Residuals | 61 | 4.6641 | 0.07646 |  | 0.30167 |  |  |
| Total | 85 | 15.4608 |  |  | 1 |  |  |
| **6 months** |  | ESV |  |  |  |  |  |
|  | Df | SumsOfSqs | MeanSqs | F.Model | R2 | Pr(>F) |  |
| Site | 4 | 5.7313 | 1.43282 | 12.4584 | 0.27056 | 1.00E-04 | *** |
| Inoculum | 4 | 4.0616 | 1.01539 | 8.8288 | 0.19174 | 1.00E-04 | *** |
| Site:Inoculum | 16 | 4.3744 | 0.2734 | 2.3772 | 0.20651 | 1.00E-04 | *** |
| Residuals | 61 | 7.0155 | 0.11501 |  | 0.33119 |  |  |
| Total | 85 | 21.1828 |  |  | 1 |  |  |
| **12 months** |  | OTU |  |  |  |  |  |
|  | Df | SumsOfSqs | MeanSqs | F.Model | R2 | Pr(>F) |  |
| Site | 4 | 6.8983 | 1.72459 | 19.7618 | 0.35664 | 1.00E-04 | *** |
| Inoculum | 4 | 2.9448 | 0.7362 | 8.436 | 0.15224 | 1.00E-04 | *** |
| Site:Inoculum | 16 | 3.216 | 0.201 | 2.3032 | 0.16627 | 1.00E-04 | *** |
| Residuals | 72 | 6.2834 | 0.08727 |  | 0.32485 |  |  |
| Total | 96 | 19.3425 |  |  | 1 |  |  |
| **12 months** |  | ESV |  |  |  |  |  |
|  | Df | SumsOfSqs | MeanSqs | F.Model | R2 | Pr(>F) |  |
| Site | 4 | 8.0058 | 2.00144 | 15.9659 | 0.31394 | 1.00E-04 | *** |
| Inoculum | 4 | 3.9817 | 0.99542 | 7.9406 | 0.15614 | 1.00E-04 | *** |
| Site:Inoculum | 16 | 4.4875 | 0.28047 | 2.2374 | 0.17598 | 1.00E-04 | *** |
| Residuals | 72 | 9.0257 | 0.12536 |  | 0.35394 |  |  |
| Total | 96 | 25.5007 |  |  | 1 |  |  |
| **18 months** |  | OTU |  |  |  |  |  |
|  | Df | SumsOfSqs | MeanSqs | F.Model | R2 | Pr(>F) |  |
| Site | 4 | 8.8405 | 2.21013 | 28.6693 | 0.4675 | 1.00E-04 | *** |
| Inoculum | 4 | 1.9093 | 0.47732 | 6.1916 | 0.10096 | 1.00E-04 | *** |
| Site:Inoculum | 16 | 2.6101 | 0.16313 | 2.1161 | 0.13802 | 1.00E-04 | *** |
| Residuals | 72 | 5.5505 | 0.07709 |  | 0.29352 |  |  |
| Total | 96 | 18.9104 |  |  | 1 |  |  |
| **18 months** |  | ESV |  |  |  |  |  |
|  | Df | SumsOfSqs | MeanSqs | F.Model | R2 | Pr(>F) |  |
| Site | 4 | 10.228 | 2.55701 | 21.3118 | 0.38457 | 1.00E-04 | *** |
| Inoculum | 4 | 3.3904 | 0.84761 | 7.0645 | 0.12748 | 1.00E-04 | *** |
| Site:Inoculum | 16 | 4.3388 | 0.27117 | 2.2601 | 0.16314 | 1.00E-04 | *** |
| Residuals | 72 | 8.6386 | 0.11998 |  | 0.32481 |  |  |
| Total | 96 | 26.5958 |  |  | 1 |  |  |

**Table S5.** PERMANOVA results for fungal OTUs vs ESVs, testing for effects of Site, Inoculum, and their interaction on Bray-Curtis community dissimilarity (data plotted in Figure S1C,D).

| **6 months** |  | OTU |  |  |  |  |  |
| --- | --- | --- | --- | --- | --- | --- | --- |
|  | Df | SumsOfSqs | MeanSqs | F.Model | R2 | Pr(>F) |  |
| Site | 4 | 3.6505 | 0.9126 | 16.292 | 0.13819 | 1.00E-04 | *** |
| Inoculum | 4 | 15.1216 | 3.7804 | 67.486 | 0.57242 | 1.00E-04 | *** |
| Site:Inoculum | 16 | 3.8354 | 0.2397 | 4.279 | 0.14519 | 1.00E-04 | *** |
| Residuals | 68 | 3.8092 | 0.056 |  | 0.1442 |  |  |
| Total | 92 | 26.4167 |  |  | 1 |  |  |
| **6 months** |  | ESV |  |  |  |  |  |
|  | Df | SumsOfSqs | MeanSqs | F.Model | R2 | Pr(>F) |  |
| Site | 4 | 3.4677 | 0.8669 | 14.285 | 0.11983 | 1.00E-04 | *** |
| Inoculum | 4 | 17.1383 | 4.2846 | 70.599 | 0.59222 | 1.00E-04 | *** |
| Site:Inoculum | 16 | 4.2065 | 0.2629 | 4.332 | 0.14536 | 1.00E-04 | *** |
| Residuals | 68 | 4.1268 | 0.0607 |  | 0.1426 |  |  |
| Total | 92 | 28.9393 |  |  | 1 |  |  |
| **12 months** |  | OTU |  |  |  |  |  |
|  | Df | SumsOfSqs | MeanSqs | F.Model | R2 | Pr(>F) |  |
| Site | 4 | 3.8139 | 0.95347 | 10.747 | 0.15252 | 1.00E-04 | *** |
| Inoculum | 4 | 11.3907 | 2.84767 | 32.096 | 0.45552 | 1.00E-04 | *** |
| Site:Inoculum | 16 | 4.123 | 0.25769 | 2.904 | 0.16488 | 1.00E-04 | *** |
| Residuals | 64 | 5.6782 | 0.08872 |  | 0.22708 |  |  |
| Total | 88 | 25.0058 |  |  | 1 |  |  |
| **12 months** |  | ESV |  |  |  |  |  |
|  | Df | SumsOfSqs | MeanSqs | F.Model | R2 | Pr(>F) |  |
| Site | 4 | 3.6894 | 0.9223 | 9.676 | 0.13472 | 1.00E-04 | *** |
| Inoculum | 4 | 12.9589 | 3.2397 | 33.987 | 0.47322 | 1.00E-04 | *** |
| Site:Inoculum | 16 | 4.6356 | 0.2897 | 3.039 | 0.16928 | 1.00E-04 | *** |
| Residuals | 64 | 6.1006 | 0.0953 |  | 0.22278 |  |  |
| Total | 88 | 27.3845 |  |  | 1 |  |  |
| **18 months** |  | OTU |  |  |  |  |  |
|  | Df | SumsOfSqs | MeanSqs | F.Model | R2 | Pr(>F) |  |
| Site | 4 | 5.111 | 1.27775 | 15.693 | 0.20105 | 1.00E-04 | *** |
| Inoculum | 4 | 10.498 | 2.62451 | 32.234 | 0.41296 | 1.00E-04 | *** |
| Site:Inoculum | 16 | 4.5197 | 0.28248 | 3.469 | 0.17779 | 1.00E-04 | *** |
| Residuals | 65 | 5.2924 | 0.08142 |  | 0.20819 |  |  |
| Total | 89 | 25.4212 |  |  | 1 |  |  |
| **18 months** |  | ESV |  |  |  |  |  |
|  | Df | SumsOfSqs | MeanSqs | F.Model | R2 | Pr(>F) |  |
| Site | 4 | 4.7856 | 1.19641 | 13.118 | 0.17019 | 1.00E-04 | *** |
| Inoculum | 4 | 12.1426 | 3.03564 | 33.284 | 0.43183 | 1.00E-04 | *** |
| Site:Inoculum | 16 | 5.2625 | 0.32891 | 3.606 | 0.18715 | 1.00E-04 | *** |
| Residuals | 65 | 5.9283 | 0.0912 |  | 0.21083 |  |  |
| Total | 89 | 28.119 |  |  | 1 |  |  |

## References

1. Frostegard A, Courtois S, Ramisse V, Clerc S, Bernillon D, Le Gall F, Jeannin P, Nesme X, Simonet P. 1999. Quantification of bias related to the extraction of DNA directly from soils. Applied and Environmental Microbiology 65:5409–5420.

2. Claesson MJ, Wang QO, O’Sullivan O, Greene-Diniz R, Cole JR, Ross RP, O’Toole PW. 2010. Comparison of two next-generation sequencing technologies for resolving highly complex microbiota composition using tandem variable 16S rRNA gene regions. Nucleic Acids Research 38.

3. Caporaso JG, Kuczynski J, Stombaugh J, Bittinger K, Bushman FD, Costello EK, Fierer N, Pena AG, Goodrich JK, Gordon JI, Huttley GA, Kelley ST, Knights D, Koenig JE, Ley RE, Lozupone CA, McDonald D, Muegge BD, Pirrung M, Reeder J, Sevinsky JR, Tumbaugh PJ, Walters WA, Widmann J, Yatsunenko T, Zaneveld J, Knight R. 2010. QIIME allows analysis of high-throughput community sequencing data. Nature Methods 7:335–336.

4. Edgar RC. 2013. UPARSE: highly accurate OTU sequences from microbial amplicon reads. Nature Methods 10:996–998.

5. Schloss PD, Westcott SL, Ryabin T, Hall JR, Hartmann M, Hollister EB, Lesniewski RA, Oakley BB, Parks DH, Robinson CJ, Sahl JW, Stres B, Thallinger GG, Van Horn DJ, Weber CF. 2009. Introducing mothur: Open-Source, Platform-Independent, Community-Supported Software for Describing and Comparing Microbial Communities. Applied and Environmental Microbiology 75:7537–7541.

6. Thompson LR, Sanders JG, McDonald D, Amir A, Ladau J, Locey KJ, Prill RJ, Tripathi A, Gibbons SM, Ackermann G, Navas-Molina JA, Janssen S, Kopylova E, Vazquez-Baeza Y, Gonzalez A, Morton JT, Mirarab S, Xu ZZ, Jiang LJ, Haroon MF, Kanbar J, Zhu QJ, Song SJ, Kosciolek T, Bokulich NA, Lefler J, Brislawn CJ, Humphrey G, Owens SM, Hampton-Marcell J, Berg-Lyons D, McKenzie V, Fierer N, Fuhrman JA, Clauset A, Stevens RL, Shade A, Pollard KS, Goodwin KD, Jansson JK, Gilbert JA, Knight R, Earth Microbiome Project C. 2017. A communal catalogue reveals Earth’s multiscale microbial diversity. Nature 551:457-+.

7. Delgado-Baquerizo M, Oliverio AM, Brewer TE, Benavent-González A, Eldridge DJ, Bardgett RD, Maestre FT, Singh BK, Fierer N. 2018. A global atlas of the dominant bacteria found in soil. Science 359:320–325.

8. Moyer CL, Dobbs FC, Karl DM. 1994. Estimation of diversity and community structure through restricition-fragment-length-polymorphism distribution analysis of bacterial 16 Sribosomal-RNA genes from a microbial mat at an active, hydrothermal vent system, Loihi seamount, Hawaii. Applied and Environmental Microbiology 60:871–879.

9. Stackebrandt E, Goebel BM. 1994. A place for DNA-DNA reassociation and 16S ribosomal-RNA sequence-analysis in teh present species definition in bacteriology. International Journal of Systematic Bacteriology 44:846–849.

10. Kunin V, Engelbrektson A, Ochman H, Hugenholtz P. 2010. Wrinkles in the rare biosphere: pyrosequencing errors can lead to artificial inflation of diversity estimates. Environmental Microbiology 12:118–123.

11. Acinas SG, Sarma-Rupavtarm R, Klepac-Ceraj V, Polz MF. 2005. PCR-induced sequence artifacts and bias: Insights from comparison of two 16S rRNA clone libraries constructed from the same sample. Applied and Environmental Microbiology 71:8966–8969.

12. Callahan BJ, McMurdie PJ, Holmes SP. 2017. Exact sequence variants should replace operational taxonomic units in marker-gene data analysis. Isme Journal 11:2639–2643.

13. Edgar RC. 2016. UNOISE2: Improved error-correction for Illumina 16S and ITS amplicon reads. doi:http://dx.doi.org/10.1101/081257.

14. Callahan BJ, McMurdie PJ, Rosen MJ, Han AW, Johnson AJA, Holmes SP. 2016. DADA2: High-resolution sample inference from Illumina amplicon data. Nature Methods 13:581-+.

15. Amir A, McDonald D, Navas-Molina JA, Kopylova E, Morton JT, Xu ZZ, Kightley EP, Thompson LR, Hyde ER, Gonzalez A, Knight R. 2017. Deblur Rapidly Resolves Single-Nucleotide Community Sequence Patterns. Msystems 2.

16. Martiny AC, Tai APK, Veneziano D, Primeau F, Chisholm SW. 2009. Taxonomic resolution, ecotypes and the biogeography of Prochlorococcus. Environmental Microbiology 11:823–832.

17. Baker NR, Allison SD. 2017. Extracellular enzyme kinetics and thermodynamics along a climate gradient in southern California. Soil Biology & Biochemistry 114:82–92.

18. Needham DM, Sachdeva R, Fuhrman JA. 2017. Ecological dynamics and co-occurrence among marine phytoplankton, bacteria and myoviruses shows microdiversity matters. ISME J 11:1614–1629.

19. Eren AM, Morrison HG, Lescault PJ, Reveillaud J, Vineis JH, Sogin ML. 2015. Minimum entropy decomposition: Unsupervised oligotyping for sensitive partitioning of high-throughput marker gene sequences. Isme Journal 9:968–979.

20. Chase AB, Karaoz U, Brodie EL, Gomez-Lunar Z, Martiny AC, Martiny JBH. 2017. Microdiversity of an Abundant Terrestrial Bacterium Encompasses Extensive Variation in Ecologically Relevant Traits. Mbio 8.

21. Dettman JR, Jacobson DJ, Taylor JW. 2006. Multilocus sequence data reveal extensive phylogenetic species diversity within the Neurospora discreta complex. Mycologia 98:436–46.

22. Chase AB, Arevalo P, Polz MF, Berlemont R, Martiny JBH. 2016. Evidence for Ecological Flexibility in the Cosmopolitan Genus Curtobacterium. Frontiers in Microbiology 7.

23. Larkin AA, Martiny AC. 2017. Microdiversity shapes the traits, niche space, and biogeography of microbial taxa. Environmental Microbiology Reports 9:55–70.

## References

1. Allison SD, Lu Y, Weihe C, Goulden ML, Martiny AC, Treseder KK, Martiny JBH. 2013. Microbial abundance and composition influence litter decomposition response to environmental change. Ecology 94:714–725.

2. Caporaso JG, Lauber CL, Walters WA, Berg-Lyons D, Huntley J, Fierer N, Owens SM, Betley J, Fraser L, Bauer M, Gormley N, Gilbert JA, Smith G, Knight R. 2012. Ultra-high-throughput microbial community analysis on the Illumina HiSeq and MiSeq platforms. Isme Journal 6:1621–1624.

3. Apprill A, McNally S, Parsons R, Weber L. 2015. Minor revision to V4 region SSU rRNA 806R gene primer greatly increases detection of SAR11 bacterioplankton. Aquatic Microbial Ecology 75:129–137.

4. Ihrmark K, Bodeker ITM, Cruz-Martinez K, Friberg H, Kubartova A, Schenck J, Strid Y, Stenlid J, Brandstrom-Durling M, Clemmensen KE, Lindahl BD. 2012. New primers to amplify the fungal ITS2 region - evaluation by 454-sequencing of artificial and natural communities. Fems Microbiology Ecology 82:666–677.

5. Looby CI, Maltz MR, Treseder KK. 2016. Belowground responses to elevation in a changing cloud forest. Ecology and Evolution 6:1996–2009.

6. Tremblay J, Singh K, Fern A, Kirton ES, He SM, Woyke T, Lee J, Chen F, Dangl JL, Tringe SG. 2015. Primer and platform effects on 16S rRNA tag sequencing. Frontiers in Microbiology 6.

7. Edgar RC. 2013. UPARSE: highly accurate OTU sequences from microbial amplicon reads. Nature Methods 10:996–998.

8. Caporaso JG, Kuczynski J, Stombaugh J, Bittinger K, Bushman FD, Costello EK, Fierer N, Pena AG, Goodrich JK, Gordon JI, Huttley GA, Kelley ST, Knights D, Koenig JE, Ley RE, Lozupone CA, McDonald D, Muegge BD, Pirrung M, Reeder J, Sevinsky JR, Tumbaugh PJ, Walters WA, Widmann J, Yatsunenkoxs T, Zaneveld J, Knight R. 2010. QIIME allows analysis of high-throughput community sequencing data. Nature Methods 7:335–336.

9. DeSantis TZ, Hugenholtz P, Larsen N, Rojas M, Brodie EL, Keller K, Huber T, Dalevi D, Hu P, Andersen GL. 2006. Greengenes, a chimera-checked 16S rRNA gene database and workbench compatible with ARB. Applied and Environmental Microbiology 72:5069–5072.

10. Koljalg U, Larsson KH, Abarenkov K, Nilsson RH, Alexander IJ, Eberhardt U, Erland S, Hoiland K, Kjoller R, Larsson E, Pennanen T, Sen R, Taylor AFS, Tedersoo L, Vralstad T, Ursing BM. 2005. UNITE: a database providing web-based methods for the molecular identification of ectomycorrhizal fungi. New Phytologist 166:1063–1068.

11. R Core Team. 2017. R: A Language and Environment for Statistical Computing, R Foundation for Statistical Computing, Vienna, Austria. http://www.R-project.org.

12. Wickham H. 2009. ggplot2: Elegant Graphics for Data Analysis. Springer-Verlag New York.

13. Oksanen J, Blanchet F, Kindt R, Legendre P, Minchin P, O’Hara R, Simpson G, Solymos P, Stevens M, Wagner H. 2012. vegan: Community Ecology Package. R package version 2.0-10.

